# Development and Validation of LAMP Primer Sets for Rapid and Correct Identification of *Aspergillus fumigatus* Carrying the *cyp51A* TR_46_ Azole Resistance Gene

**DOI:** 10.1101/2021.03.02.433685

**Authors:** Plinio Trabasso, Tetsuhiro Matsuzawa, Teppei Arai, Daisuke Hagiwara, Yuzuru Mikami, Maria Luiza Moretti, Akira Watanabe

## Abstract

Infections due to triazole resistant *Aspergillus fumigatus* are increasingly reported worldwide and are associated with treatment failure and mortality. The principal class of azole resistant isolates is characterized by the presence of tandem repeats of 34 bp or 46 bp (TR_34_ or TR_46_) within the promoter region of the *cyp51A* gene. Loop-mediated isothermal amplification (LAMP) is a widely used nucleic acid amplification system with high rapidity and specificity. In this paper, we report a new LAMP assay method to detect the 46 bp tandem repeat insertion in the *cyp51A* gene promoter region, named TR_46_-LAMP assay, based on the use of a newly designed specific LAMP primer sets. TR_46_ is a high-prevalence allele which is associated with the occurrence of multi-triazole resistance of *A. fumigatus* in patients as well as isolates from the environment. This newly designed TR_46_-LAMP assay was validated as a useful method for specific detection of azole-resistant *A. fumigatus* isolates bearing TR_46_^2^ as well as TR_46_^3^ in *cyp51A* gene promoter region. It could also differentiate azole-resistant isolates of TR_46_ tandem repeats from those with TR_34_ tandem repeats in *cyp51A* genes. These results showed this TR_46_-LAMP method is specific, rapid, and also provides crucial insights to enable development of novel antifungal therapeutic strategies against severe fungal infections due to *A. fumigatus* with TR_46_ tandem repeats.

## INTRODUCTION

Antimicrobial resistance was defined by the World Health Organization (WHO) as one of the most important threats to human health, since it can compromise our ability to treat infectious diseases, as well as undermining other advances in health care (1). Although bacterial resistance still remains the most common finding in the clinical setting, fungal resistance, specially to azole antifungals among filamentous fungi, is relentlessly increasing worldwide (2,3,4). Resistance to azole drugs can be due, in general, to two major genetic mechanisms; point mutation(s) in *cyp51A* open reading flame with or without tandem repeat (TR) of 34 or 46 base pair (bp) in the promoter region of the gene (TR_34_ or TR_46_) and overexpression of oligonucelotide sequence in the cyp51A gene (5). These mutations and overexpression of the gene are conferring different levels of resistance (6). As a general concept, the point mutation is the consequence of previous exposure to azole drugs in the clinical setting, such as for prophylactic approaches or for therapeutic purpose (7,8). On the other hand, the TR with point mutation(s) is understood as the aftermath of previous exposure to azole fungicides. In an agricultural setting, azole fungicides are widely used to prevent fungal contamination in a large variety of crop and plant protection, which may allow the TR type resistant strains to be emerged in the environments and the conidia to disperse to the air (8,9). In Brazil, agrobusiness represents about 25% of the Brazilian Gross National Product (GNP) (http://www.agricultura.gov.br), and in this scenario, fungicides use is steadily increasing over the years. In 2010, the consumption of pesticides in Brazil increased 190% in the year and fungicides corresponded to 14% in this market (https://www.ipessp.edu.br).

In the Netherlands, prevalence of TR type resistant strains were seriously high. A set of 952 clinical *A. fumigatus* strains were collected and found to include 225 and 98 strains with TR_34_ and TR_46_, respectively (10). Another study revealed that 13 TR_34_ strains and 3 TR_46_ strains were isolated among 105 clinical strains. Accordingly, there are number of reports assessing fatal case of invasive aspergillosis caused by *A. fumigatus* carrying the TR_46_ in an acute myeloid leukemia (AML) patient and hematopoietic stem cell transplant recipients (11). Higher mortality of the patients with invasive aspergillosis caused by azole resistant strain has been reported (11,12). Thus, a rapid and specific method which is able to identify the presence of TR would contribute to more timely therapeutic decision making (13).

As one of the promising diagnostic tools for the azole resistant *A. fumigatus*, the high potential of loop-mediated isothermal amplification (LAMP) for the development of improved DNA-based diagnostic kits has been reported (14). In general, the LAMP method was found to be similar or superior to the general PCR method, and more specific, low-cost and simple. Actually LAMP based approaches have been applied to a wide range of samples, such as whole blood, paraffin-embedded tissues, and various microbial pathogens. In this paper, we report a novel LAMP assay method which selectively detects triazole resistant *A. fumigatus* strains due to presence of double or triple TR_46_(TR_46_^3^) in *cyp51A* promoter region.

## RESULTS

### Antifungal susceptibility tests

Drug susceptibilities of 41 *A. fumigatus* strains against azole drugs itraconazole and voriconazole are shown in Table 2. Thirty strains designated as wild type, were isolates from clinical specimens, and they were confirmed to belong to drug susceptible to itraconazole and voriconazole. Remaining 11 strains were in the azole resistant group, and most of them showed MIC values of >8*μ*g/ml against voriconazole.

### Primer design

The most important step in LAMP assay is the design of primers. In LAMP assay, 6 primers are necessary to amplify the targeted region under isothermal condition. First, we inspected the promoter region (−461 bp to – 296 bp counted from start codon) of *cyp51A* gene to select a set of primer sequences that specifically amplify the repeated 46 bp sequence in strains with TR_46_ mutation (Fig. 1). To enable for specific amplification against repeated TR_46_ sequences, B2 was set on the joint with two 46 bp sequences. To obtain a specific and rapid LAMP primer set in the LAMP assay, other 5 sequences for primer sets were chosen in the target region according to the standard criteria. Namely 6 primers (F1, F2, F3, B1, B2, B3) that target 6 specific regions of a DNA template of the TR_46_ gene of *cyp51A* were selected, and in addition 2 loop primers (LF, LB) were also chosen to accelerate the reaction. Based on the above information, several new candidates of LAMP primers were designed and their utility was tested. From those, one useful LAMP primer sets based on the detection of TR_46_ regions in *cyp51A* gene was selected (Table 1).

**Fig. 1.**
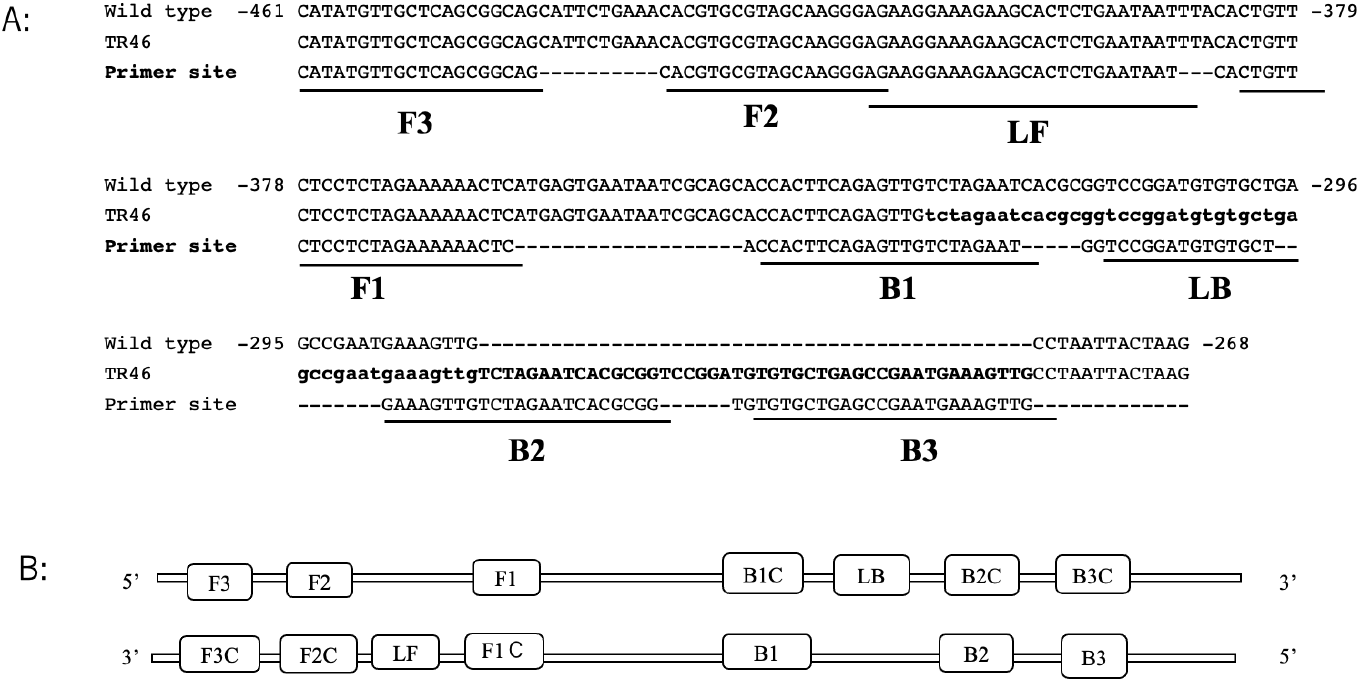
Genetic information for the design of the LAMP primer set sequence. A: Schematic illustration of *cyp51A* gene showing LAMP primer positions and corresponding sequences of TR_46_ bp promoter tandem repeat sequences in comparion with those of wild type. B: Primers F3, F2, F1, B1, B2 and B3 show primer sequence positions. Sequences of some primers are complementary as shown in Table 1. See LAMP primer and methods which are shown in refereces 19 and 20.

**Table 1.**
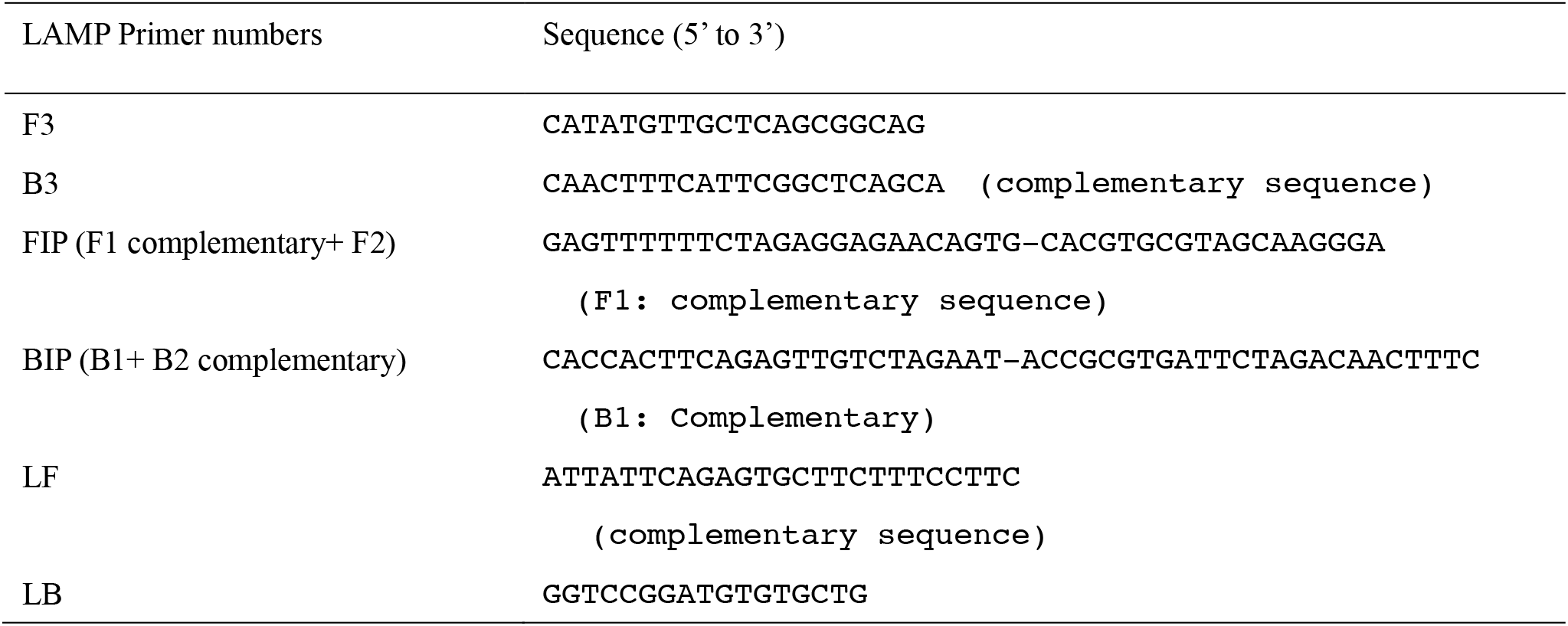
Sequence information of newly designed and useful TR_46_-LAMP primers set in this experiment

### Verification of LAMP for TR_46_

Specificity of the primer sets was tested using various types of *A. fumigatus* strains, such as reference wild isolates, and environmental or clinical azole resistance isolates (Fig. 2).

**Fig. 2.**
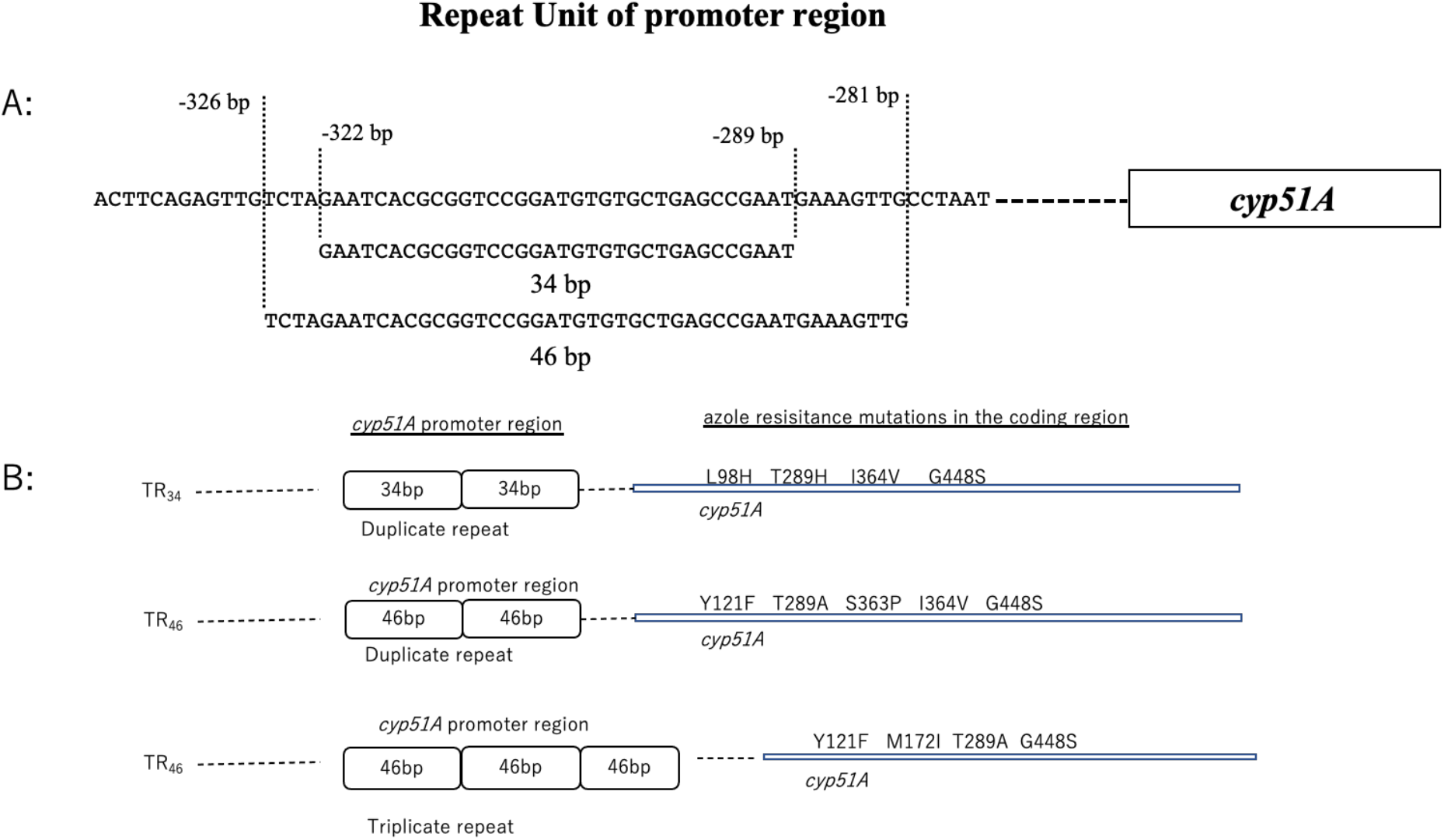
Illustration of tandem repeat regions of *cyp51A* genes used in this experiment. A: Tandem repeat unit of promoter genes of TR_34_ and TR _46_. B: Tandem repeat: 34 bp (double) and 46 bp (double and triple), and *cyp51A* gene associated point mutation place.

As shown in Fig. 3, the start of the LAMP amplification was at around 50 min in the *A. fumigatus* strains (IFM 63432) harboring the TR_46_ resistant mutation (TR_46_/Y121F/T289A). In the present study, IFM 63432 strain was used as a positive control one for this LAMP amplification experiment. This TR_46_ LAMP primer could not amplify the DNA from 30 strains of azole drug susceptible wild type and clinical isolates of *A. fumigatus* (Fig. 3 A-i). As shown in Fig. 3A-ii, TR_46_ LAMP primer could amplfy the DNA from *A. fumigatus* strains carrying the duplicate 46 bp promoter repeat in *cyp51A* gene (IFM63432, Be1-2, BE1-4, BE3-5, BE3-6). When the present TR_46_ LAMP primer was applied for three strains with TR_34_ (Table2), TR_34_/L98H (IFM64460 and 64733) and TR_34_/L98H/Y289/T289A/I364V/G448S (3-1-B), DNA amplification was not observed. This suggests that the present LAMP primer could not detect TR_34_ drug resistant strains regardless of their point mutation site in *cyp51A* gene (Fig. 3 A-iii and B-ii). It was also confirmed that TR_46_ LAMP primer could amplify DNA of *A. fumigatus* strains carrying a TR_46_^3^ (TR_46_^3^/Y121F/M172I/T289A/G448S) in the *cyp51A* gene (Fig.3A-iii,3B-ii). Therefore, present studies confirmed that newly established TR_46_ LAMP primer set was only specific *A. fumigatus* strains with TR of double or triple 46-bp promoter tandem repeats in *cyp51A* gene. Colony LAMP verified for 2 strains each carrying TR_46_, TR_46_^3^, and TR_34_.

**Fig. 3.**
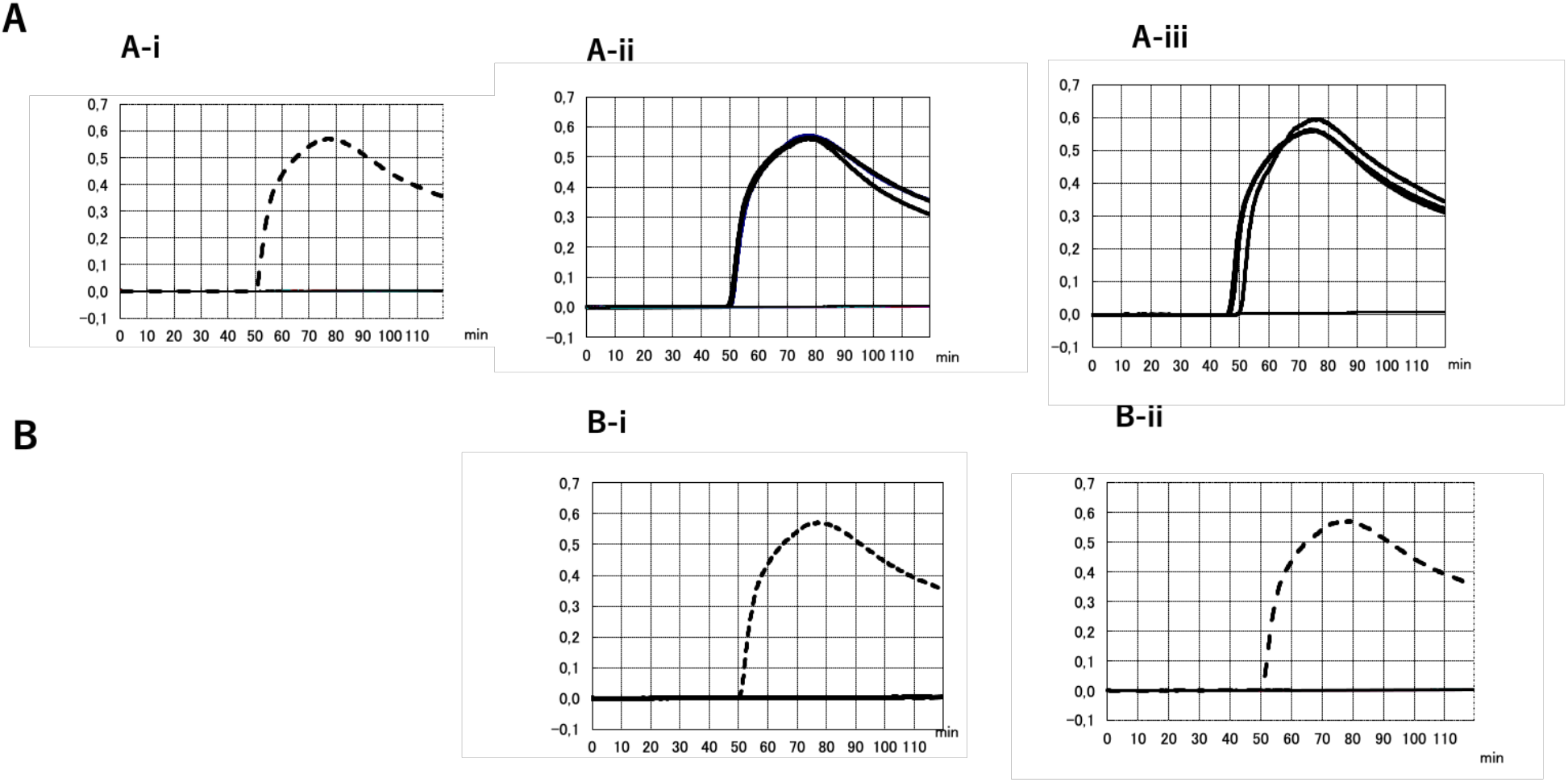
Comparative amplification profiles of *A. fumigatus* wild type isolates, and environmental or clinical azole resistant isolates with or without TR_46_ double or triple 46 bp promoter repeats in *cyp51A* gene by a newly developed LAMP primer set. Amplification curve of positive control strain (IFM63432) is shown by dotted line. A-i:DNA amplification profiles using 30 strains of *A. fumigatus* wild type. DNA amplification was not comfirmed in the wild type strains tested (30 strains). A-ii: DNA amplification was comfirmed by four TR_46_^2^ strains (IFM 63432, BE1-2, BE1-4, BE3-5, BE3-6) which have double 46 bp promoter repeats. A-iii: DNA amplification was confirmed by three TR_46_^3^ strains (BE1-1, W1-4, W2-12-1: Y121F/M1721I/T289A/G448S) which have triple 46 bp promoter repeats. IFM63432 strain was used as a positive control. B-I: DNA amplification was not confirmed by three TR_46_^3^ strains (BE1-1, W1-4, W212-1) which have duplicate 34 bp promoter repeats and one mutations in the coding region(L98H). B-ii: DNA amplification was not confirmed by one TR_34_^2^ strain (3-1-B) which has duplicate 34 bp promoter repeats and multi-mutions in the coding region (L98H/T289A/I364V/G448S).

**Table 2.** *Aspergillus fumigatus* strains used in this experiment

| Strain No. | Isolation source | *cyp51A* genotypes | MIC values ($\mu$ g/ml) | |
| --- | --- | --- | --- | --- |
|  |  |  | ITCZ | VRCZ |
| IFM63432 | clinical | TR<sub>46</sub>/Y121F/T289A | 4 | >8 |
| BE1-2 | environmental (bulb)<sup>a</sup> | TR<sub>46</sub>/Y121F/T289A | 2 | >8 |
| BE1-4 | environmental (bulb) | TR<sub>46</sub>/Y121F/S363P/I364V/G448S | 2 | >8 |
| BE 3-5 | environmental (bulb) | TR<sub>46</sub>/Y121F/T289A | 2 | >8 |
| BE 3-6 | environmental (bulb) | TR<sub>46</sub>/Y121F/T289A | 2 | >8 |
| BE 1-1 | environment (bulb) | TR<sub>46</sub><sup>3</sup>/Y121F/M172I/T289A/G448S | 2 | >8 |
| W1-4 | environment (bulb) | TR<sub>46</sub><sup>3</sup>/Y121F/M172I/T289A/G448S | 2 | >8 |
| W2-12-1 | environment (bulb) | TR<sub>46</sub><sup>3</sup>/Y121F/M172I/T289A/G448S | 2 | >8 |
| IFM64460 | clinical | TR<sub>34</sub>/L98H | >8 | >8 |
| IFM64733 | environmental | TR<sub>34</sub>/L98H | >8 | >8 |
| 3-1-B | environment (bulb) | TR<sub>34</sub>/L98H/T289A/I364V/G448S | 2 | >8 |
| IFM62918 | clinical | wild | 0.5 | 1 |
| IFM62799 | clinical | wild | 0.5 | 1 |
| IFM60516 | clinical | wild | 1 | 1 |
| IFM58402 | clinical | wild | 0.5 | 0.5 |
| IFM51977 | clinical | wild | 0.25 | 0.25 |
| IFM60065 | clinical | wild | 1 | 0.5 |
| IFM61960 | clinical | wild | 0.5 | 0.5 |
| IFM51748 | clinical | wild | 0.125 | 0.125 |
| IFM63666 | clinical | wild | 1 | 2 |
| IFM62520 | clinical | wild | 1 | 0.5 |
| IFM50999 | clinical | wild | 0.5 | 0.5 |
| IFM50268 | clinical | wild | 0.25 | 0.125 |
| IFM55548 | clinical | wild | 0.25 | 0.25 |
| IFM63311 | clinical | wild | 1 | 0.5 |
| IFM63355 | clinical | wild | 2 | 2 |
| IFM60901 | clinical | wild | 0.5 | 0.5 |
| IFM62674 | clinical | wild | 1 | 2 |
| IFM62709 | clinical | wild | 0.5 | 0.5 |
| IFM52659 | clinical | wild | 1 | 1 |
| IFM57130 | clinical | wild | 0.25 | 0.125 |
| IFM60814 | clinical | wild | 0.5 | 0.5 |
| IFM49435 | clinical | wild | 0.25 | 0.25 |
| IFM61572 | clinical | wild | 0.5 | 0.5 |
| IFM50669 | clinical | wild | 0.5 | 0.25 |
| IFM59832 | clinical | wild | 0.5 | 0.5 |
| IFM55044 | clinical | wild | 0.25 | 0.25 |
| IFM47670 | clinical | wild | 0.5 | 0.5 |
| IFM51978 | clinical | wild | 0.5 | 0.25 |
| IFM58328 | clinical | wild | 0.5 | 0.5 |
| IFM60369 | clinical | wild | 0.5 | 0.5 |

a: obtained from plant bulbs.

## DISCUSSION

Azole antifungals mainly inhibit ergosterol biosynthetic pathway by targeting the cytochrome P450-dependent enzyme lanosterol 14-α-demethylase, encoded by *cyp51A* in molds. Resistance to this class of drugs in the major human pathogen *A. fumigatus* is emerging and reaching levels to prevent their clinical use (6). Advances in recent molecular genetic technologies such as real-time PCR have introduced various useful diagnostic assay methods into the fields in azole resistant mechanism analysis. Even in comparison with those of new molecular biological techniques, the LAMP assay described here has advantages of high sensitivity and specificity, and also low costs and short amplification time. In addition, there have been no reports using LAMP techniques to study azole resistant mechanisms in *A. fumigatus* by TR_46_ gene. Quite recently Yu Shan-Ling *et al.* (15) reported a successful LAMP technique to detect azole resistant strains due to amplification of a TR of a 34 bp (TR_34_) within the promoter region of *cyp51A* of *A. fumigatus.* However, their method could not detect azole resistant *A. fumigatus* strains with amplification of a 46 bp of promoter regions. There is a difference in MIC between strains with TR_46_ and strains with TR_34_ (16). Thus, the importance of detecting TR_46_ lies in the fact that strains of *A. fumigatus* harbouring TR_46_ are resistant to voriconazole, but not to itraconazole, while strains with TR_34_ are resistant to both drugs (16). In this study, we were succeeded in the design of useful TR_46_ LAMP primer sets to detect specifically a TR_46_ within the promoter regions of azole resistant *A. fumigatus*. The designed primer sets could differentiate azole resistant TR_46_ strains from that of TR_34_ strains. To our knowledge, this is the first report of a detection method for one of the most prevalent *cyp51A* resistant gene TR_46_ in *A. fumigatus* azole resistant strains.

Recently, the strain consisting of the triple repeat of 46 bp of the promoter region was reported in the Netherlands (TR_46_^3^) (17). The LAMP primer we designed was able to detect both two copies of the TR_46_ tandem repeat and three copies of the TR_46_. Moreover, these amplification curves (as well as the starting point) were similar. The BIP (B1 + B2 complementary; Fig. 1) of the primer we designed is TR-specific. B1 is designed at the boundary where the repeat unit is inserted, and B2 is designed at the boundary between the repeat units. In addition, B3 is designed on a repeat unit. Based on the results of strains having double repeat and triple repeat, it was suggested that the primer used this time may be able to detect even if the number of repeats increases.

It is widely known that exposure to azole fungicides resulted in the emergence of azole-resistant strains with tandem repeats in the promoter region of cyp51A. (8,9) For this reason, epidemiological studies such as the incidence of azole-resistant strains in the environment are important. Many environmental and clinical isolates need to be screened to generate epidemiological data such as the frequency of detection of azole-resistant *A. fumigatus*. The method developed in this study would be an easy-to-use screening procedure.

Since the LAMP assay developed in the present study is one-step and rapid detection method, coupled with its high reliability and ease of use, it can be used for prompt specific detection of drug resistant genes due to TR_46_ in *A. fumigatus* in the clinical laboratory. Early detection of infections due to TR_46_ drug resistant strains in *A. fumigatus,* might be helpful to guide the early start of corrective and effective antifungal therapy.

## MATERIALS AND METHODS

### *Aspergillus* isolates

Forty-one strains, including thirty-three from the clinical setting and eight environmental (plant bulbs) isolates (18) of *A. fumigatus* were provided through the National Bio-Resource Project (NBRP), Japan (http://www.nbrp.jp/); source and drug susceptibility are shown in Table 2.

### DNA preparation and extraction

These fungal strains were cultured on Sabouraud dextrose agar. Genomic DNA was extracted from over-night cultures of *A. fumigatus* mycelia by urea-phenol method. Mycelia were mixed with 0.5 mm size glass beads, 0.5 ml of PCI (phenol/chloroform/Isoamyl alcohol) solution and 0.5 ml DNA extraction buffer (50 mM Tris-HCl, pH 8.0, 20 mM EDTA, 0.3 M NaCl, 0.5% SDS, 5 M urea), and disrupted by FastPrep FP100A (MP-Biomedicals, Santa Ana, USA). After centrifugation, the upper layer was transferred to a new tube and subjected to ethanol precipitation. The resulting DNA pellet was suspended in 100 μl TE buffer. DNA concentration was determined by the methods described in our previous paper (19).

### MIC determination by broth microdilution test

All *A. fumigatus* strains were submitted to antifungal susceptibility tests according to the CLSI M38 protocol (https://clsi.org/standards/products/microbiology/documents/m38/), using Eiken Dried Plates (9DEF47, Eiken Chemical Co., Tokyo, Japan).

### LAMP-method

LAMP was performed as described in our previous studies (20). TR_46_ LAMP primer was designed base on the target promoter region sequences of *cyp51A* gene of *A. fumigatus*, which includes tandem repeats in the promoter region containing TR_46_ mutant alleles. The sequence information of *cyp51A* gene were downloaded from NCBI Gen-Bank (https://www.ncbi.nlm.nih.gov/genebank). In total, a 184-bp nucleotide alignment (Fig. 1) was used for TR_46_ LAMP primer design by the protocol reported by Eiken Company (PrimerExplorer V5, Eiken Chemical Co. Ltd, Tokyo. Japan). LAMP primers are composed of six primers recognizing six distinct regions. The forward and backward inner primers, FIP/BIP, play crucial roles in the specificity of the assay. The outer primers, F3/B3, are composed of the fewer bases and are of a lower concentration than are FIP/BIP, initiating annealing of F3/B3 to the target in order to commence strand displacement. In addition to these four essential primers (FIP/BIP and F3/B3), the forward and backward loop primers (LF/LB) are used. The high specificity and rapidity of the present LAMP assay were achieved by applying 6 primers that target 6 regions of a DNA template, and 2 loop primers (LF, LB) to accelerate the reaction. LAMP reactions were performed with a Loopamp DNA amplification kit using reaction mixtures composed of 40 pmol each of primers FIP and BIP, 5 pmol each of primers F3 and B3, 20 pmol each of primers LF and LB, 12.5 ml ×2 reaction mixture, 1 μl Bst DNA polymerase, 2 μl DNA sample and distilled water up to a final volume of 25 μl (Eiken Chemical Co., Ltd., Tokyo, Japan). The LAMP reactions were analyzed by a real-time turbidimeter(Loopamp EXIA; Eiken Chemical Co.) and were conducted at 63°C, for 120 min. The reaction mixtures were incubated at 61℃ for 30 min (Realoop-30; Eiken Chemicals, Japan), and then heated at 80℃ for 2 min to terminate the reaction. Start of amplification of LAMP products at 30 to 50 minutes in the graph, suggested the positive reaction due to the presence of corresponding a 46 bp tandem repeats of *cyp51A* gene is short. Since overall reaction can be obtained within 2 hours, prompt drug therapy can be deployed within a short time.

## ACKNOWLEDGEMENTS

This study was supported by AMED under Grant Numbers JP20jm0110015 and by JICA through the collaborative research project Science and Technology Research Partnership for Sustainable Development (SATREPS), Japan.

